# Polystyrene-colonizing bacteria are enriched for long-chain alkane degradation pathways

**DOI:** 10.1101/2022.11.18.516359

**Authors:** Shu Wei Hsueh, You-Hua Jian, Sebastian D. Fugmann, Shu Yuan Yang

## Abstract

One of the most promising strategies for the management of plastic waste is microbial biodegradation, but efficient degraders for many types of plastics are still lacking, including for polystyrene (PS). Genomics has emerged as a powerful tool for mining environmental microbes that may have the ability to degrade different types of plastics. In this study, we use 16S sequencing to analyze the microbiomes for multiple PS samples collected from sites with different vegetation in Taiwan to reveal potential common properties between species that have exhibit growth advantages on PS surfaces. Phylum enrichment analysis identified Cyanobacteria and Deinococcus-Thermus as being the most over-represented groups on PS, and both phyla include species known to reside in extreme environments and could encode unique enzymes that grant them properties suitable for colonization on PS surfaces. Investigation of functional enrichment in PS-enriched species highlighted carbon metabolic pathways, especially those related to hydrocarbon degradation. This is corroborated by the finding that genes encoding long-chain alkane hydroxylases such as AlmA are more prevalent in the genomes of PS-associated bacteria. Our analyses illustrate how plastic in the environment support the colonization of different microbes compared to surrounding soil. In addition, our results point to the possibility that alkane hydroxylases could confer growth advantages of microbes on PS.

## Introduction

It is widely accepted that the plastic waste accumulated and continuously increasing in the world presents one of the gravest environmental challenges we currently face. One potential solution is to mine the diverse world of microbes that carry a broad range of enyzmes for the ability to degrade different types of plastics. The most efficient examples known to act on plastics are those that degrade polyethylene terephthalate (PET) into monomers, such as the PETase-MHETase enzyme combination isolated from *Ideonella sakiensis* and leaf branch compost cutinase (1, 2).

For the biodegradation of polystyrene (PS), several studies have reported modest results by individual microbial strains and consortia (3–9). In 2015, larvae of yellow mealworms (*Tenebrio molitor*) were shown to ingest and degrade the extruded form of PS (EPS, or Styrofoam) (10). Since then, the larvae of several other species including dark mealworm (*Tenebrio obscurus*), superworm (*Zophobas atratus*), wax moth (*Galleria mellonella*), and beetles have also been shown to exhibit similar behavior and abilities (11–14). Mineralization of PS in *Tenebrio molitor* occurs in the gut via the help of the associated microbiota (15, 16), but few enzymes and pathways that mediate such degradation have been defined.

PS, like polyethylene (PE) and polypropylene (PP), has a carbon backbone that makes them highly refractory to biodegradation. In considering possible biological pathways that may be able to degrade such plastics, an interesting point of reference is the degradation of petroleum hydrocarbons which are similarly hydrophobic and contain a large portion of alkanes. Microbes are well-documented to perform bioremediation of oil-polluted environments by degradation of hydrocarbons (17). These processes are frequently mediated by oxidation via alkane hydroxylases (18), and the similarities between the properties of petroleum hydrocarbons and plastics make alkane hydroxylases interesting candidates for degrading plastics whose polymer backbone consists of carbon-carbon bonds. Indeed, there is accumulating evidence indicating that alkane hydroxylases such as AlkB can transform PE (19), and these enzymes have also been implicated in oxidizing PS (20–22).

To investigate the potential of bacterial enzymes for PS degradation, we surveyed the microbiomes of multiple EPS waste samples from various environmental sites by 16S amplicon sequencing. Functional analyses revealed that metabolism of hydrocarbons is enriched in PS-associated species, and subsequent examination of the presence of alkane hydroxylases revealed that the enzymes that degrade long-chain alkanes show the strongest enrichment. These observations highlight long-chain alkane hydroxylases as novel candidates of PS biodegradation.

## Methods

### Sample collection and 16S rRNA gene sequencing

EPS waste pieces were collected at 7 different sites of varying location, vegetation, and pH, and two control soil samples were also collected from each site, one directly underneath the EPS sample (“S” for soil) and another 0.5 m away (“SN” for soil nearby). The pH values for all soil samples were measured with pH paper of soil mixed with equal parts water (HYDRION acid-base test paper, Micro Essential Laboratory).

To extract environmental DNA from the EPS samples, microbes from the collected pieces were first submerged in no carbon media (26.1 mM Na_2_HPO_4_, 22 mM KH_2_PO_4_, 8.6 mM NaCl, 18.7 mM NH_4_Cl, 0.4 mM MgSO_4_, 36 mM FeSO_4_, 29.6 mM MnSO_4_, 47 mM ZnCl_2_, 4.6 mM CaCl_2_, 1.3 mM CoSO_4_, 1.4 mM CuSO_4_, 0.1 mM H_3_BO_3_, 34.2 mM EDTA, 1.8 mM HCl), vortexed for 5-10 min, and subjected to DNA extraction with the DNeasy PowerWater Sterivex kit (Qiagen) as instructed by the manual. DNA from soil samples were extracted with the DNeasy PowerSoil Pro kit (Qiagen) following manufacturer’s protocol. These DNA samples were submitted for full length 16S rRNA gene sequencing on the PACBIO SMRT platform and the average raw HiFi reads obtained per sample was 16055.

### Data analysis

The reads obtained for individual sequences were processed with DADA2 to obtain amplicon sequence variants (ASVs) which are then referenced with databases (NCBI, GreenGenes, SILVA, eHOMD, UNITE) to generate ASV tables that included taxonomy assignments. The ASV tables were then used for the determination of Simpson diversity indices and rarefaction curves to indicate alpha diversity, and for PCA analysis to calculate beta diversity. Functional enrichment analysis was performed with the FAPROTAX database (26).

To determine top enriched species associated with PS, the read numbers for each ASV was first normalized against total reads in a sample before calculating average enrichment values by dividing the average normalized reads of an ASV across PS samples with the average for either the S or SN samples (Table S3). This enabled the ranking by enrichment values of all species detected whose whole genome sequences are available to generate the top enriched species list. Control lists of species were selected randomly excluding species whose whole genome was yet unavailable or that overlapped with the top 50 enriched list (Table S3).

### Identification of alkane hydroxylases homologs

To identify homologs of alkane hydroxylases, well-characterized homologs of each type were used as the query sequence to blast against whole genome sequences (tblastn) of designated species and strains at the NCBI database and further examined for the presence of signature domains or residues if such information is available.

For AlkB, homologs were identified based on homology to AlkB from *Pseudomonas putida* (Q9WWW6) and the presence of conserved motifs including 3 histidine boxes (HELXHK, EHXXGHH, LQRHSDHHA) and an HYG motif (NYXEHYGL) (Smits 2002); those that belong to the LLM class of flavin-dependent oxidoreductase were excluded. The query sequence for identifying CYP153 homologs was from *Mycobacterium marinum* (B2HGN5).

For AlmA, the homolog from *Acinetobacter baylyi* (Q6F7T9) was used as the query. Hits that contained at least a region of 250 a.a. of 25-50% identity to the query and a predicted flavin-binding monooxygenase (FMO-like) domain or CzcO domain, both related to alkane hydroxylases, were called AlmA homologs. LadA homologs were determined similarly using the homolog from *Geobacillus thermodenitrificans* (A4IU28) as the query for tblastn, and the hits we called LadA showed at least 200 a.a. of homology to the query with 20-50% sequence identity. They also contained flavin-utilizing monooxygenenase domains.

For methane monooxygenases, protein sequences from *Thauera butanivorans* (A7MAQ9), *Nocardioides sp. CF8* (E9L6F6), *Mycobacterium sp. TY-6* (Q08KF2), and *Pseudonocardia sp. TY-7* (Q08KE2) were used as the query for BmoR, BmoA, PrmA and Prm1A, respectively.

Classification of AlmA and LadA proteins included homologs identified in the top 30 PS-enriched species along with those from the random 60 species list as we wanted to include a greater number of homologs from species that did not exhibit enrichment on PS (Table S4). Phylogenetic analyses of these homologs was carried out first by aligning the regions homologous to the AlmA and LadA queries with the ClustalW tool. Trees were subsequently built with MEGA7 using the Maximum Likelihood method and performing 1000 bootstrap replicates (44); bootstrap consensus trees are presented.

### Comparisons of multiple *almA* genes in strains of a species

There is one fully assembled genome each available for *Pseudomonas toyotomiensis* (SM2) and *Pseudomonas alcaliphila* (JAB1), and 4 and 2 additional whole genome projects for the two species, respectively (*P. toyotomiensis:* KF710, JCM 15604, DSM 26169, 718, SM2; *P. alcaliphila:* JCM 10630, NBRC 102411). For all strains, homologs of AlmA were determined by tblastn as described above and in the Results section. In addition, the two genes immediately upstream and downstream to each *almA* were identified for synteny determination.

## Results

### Profiling of PS-colonizing microbiota

To investigate the microbiomes on PS samples in natural environments, we collected 7 pieces of waste EPS from different sites in Taiwan (Fig. 1). These samples were all partially embedded in surrounding soil and showed clear signs of weathering, suggesting that these EPS pieces have been in those environments for substantial amounts of time which could allow for microbial communities to colonize and mature. If any microbes have the potential to take advantage of the plastic material, such as using it as a survival niche or an alternative carbon source, those strains would gain growth advantages and exhibit enrichment compared to in the soil in their proximity. Due to these considerations, for each EPS waste piece we also collected two control soil samples, one directly underneath the EPS piece we collected (“S” for soil), and another one 0.5 m away (“SN” for soil nearby). For each site of collection, we recorded the vegetation around where the EPS sample was collected, and the pH values of the soil samples were determined (Table S1). To analyze the bacterial microbiomes associated with the EPS and soil samples we collected, we performed 16S rRNA gene sequencing to determine the species composition and abundance for all 21 samples. We chose to sequence fulllength 16S genes to achieve greater species resolution.

**Figure 1.**
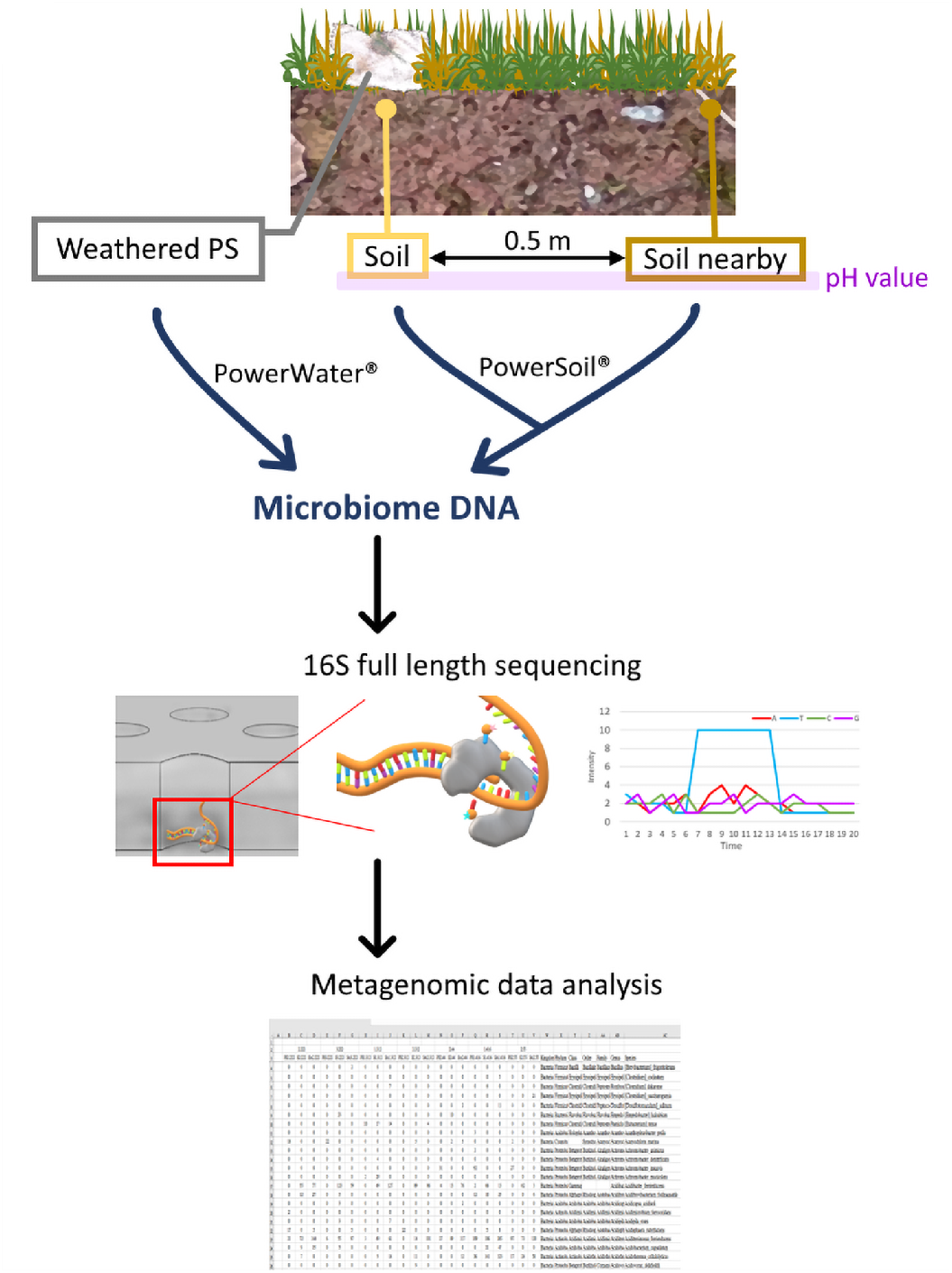
Diagram of sample collection and processing. Weathered EPS samples were collected from 7 different locations; at each location two control soil samples were also collected, one directly underneath the EPS piece (S for soil) and another 50 cm meters away (SN for soil nearby). DNA was extracted from the samples using commercial kits and subsequently submitted for full length 16S sequencing to survey the associated microbiomes.

### PS-residing microbiota is significantly different from those in surrounding soil samples

The first test we applied to our genomics data was to investigate whether there are differences in the microbiota observed on PS samples compared to the soil ones. Using PCA, we found that bacterial contents on the PS samples as a group was significantly different compared to those of either the S or SN samples (Fig. 2A). The microbial compositions on soil samples were quite similar to one another in spite of the differences in location, vegetation, and soil pH of the collection sites (Table S1); in comparison there is much more variation in the microbiomes associated with PS samples. Another finding of note is that the alpha diversities of PS-associated microbiomes were less compared to those in soil (Fig. S1A-B). This indicates that the PS surface supports colonization of a more limited repertoire of microbes, which is what some reports studying microbiomes on other petroleum-based plastics have also found (23, 24). The differences between the three types of samples can also be observed when we examined the bacteria composition based on taxonomy. The abundance levels of the most prevalent phyla and species are different on PS compared to the soil controls (Fig. 2B, S2), and the most highly enriched species on PS, S, and SN samples are also distinct as evidenced by analysis using the LEfSe method (Fig. S3) (25).

**Figure 2.**
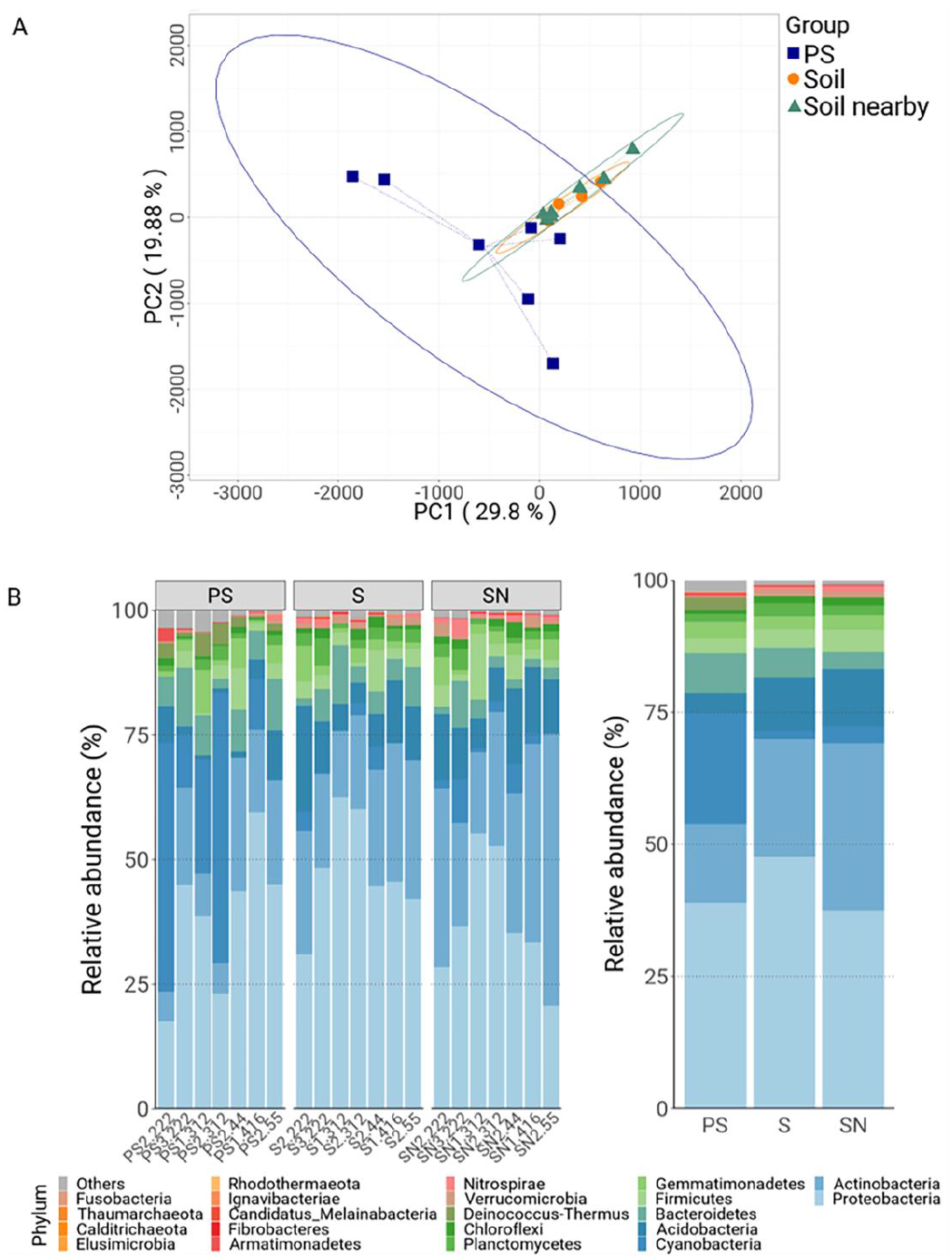
Comparisons of PS- and soil-associated microbiomes. **A,** PCA analysis of the microbiomes collected off PS (blue), S (orange), and SN (green) samples. **B,** Bar graphs showing the relative levels of the most-abundant phyla for the PS, S, and SN samples with individual specimens of each type displayed separately (left graphs) or collectively (right graph). Color codes of the different phyla are to the right of the graph.

These analysis results, taken together, indicate that the microbial community colonizing EPS is significantly different from those found in the surrounding soil environments. This is in spite of the proximity of PS samples to the soil samples and that the microbes on PS most likely having originated from their immediate environments. We think this shows that PS, due to its chemical composition or acting as a niche, is able to nurture specialized localization and growth of microbes.

### Enrichment of Bacterial Phylum on PS samples

To investigate how PS-associated microbiomes are distinct from those of the surrounding environments, we examined whether there are certain bacterial clades that are particularly abundant in the PS samples compared to the soil controls. At the phylum level, we found that the abundances of Cyanobacteria and Deinococcus-Thermus on PS samples are clearly higher than those in controls (Fig. 2B). Interestingly, the enrichment is due to multiple species from the two phyla being present and not due to the enrichment of one or two outstanding species (Table S2), suggesting that common properties of species within these two phyla underlie their ability to survive on PS. One of the best known characteristics of cyanobacteria is their ability to perform photosynthesis, and we wondered whether the prevalence of cyanobacteria on the PS samples we collected was due to them being readily exposed to sunlight. However, the SN samples we collected were also surface samples that had ample exposure to sunlight, and the PS samples showed comparable enrichment in cyanobacteria compared to both the S and SN samples. Therefore, the higher abundance of cyanobacteria on PS samples is not due to the availability of sunlight.

The more interesting idea of why cyanobacteria exhibit PS enrichment would be that they are able to benefit from PS as a source of nutrients either by directly degrading PS or utilization of PS metabolites generated by other microbes. We did not find any metabolic pathways or enzymes of note in Cyanobacteria or Deinococccus-Thermus species enriched on PS compared to controls (Fig. S4A), but this certainly does not rule out the possibility that other novel enzymes would be able to contribute to survival advantages of these phyla on PS.

### Genomics analysis reveal an enrichment of hydrocarbon degradation pathways in microbiota on PS samples

To investigate whether the bacterial community present on PS show signs of being able to utilize the plastic as a source of nutrients, we looked for enrichment of metabolic pathways of species associated with PS samples compared to surrounding soil samples. Using the FAPROTAX database (26), we tested whether any metabolic pathway is more abundantly represented in microbes present on PS, and found two types to show the clearest and highest enrichment: photosynthetic functions and carbon metabolism, especially those related to alkane metabolism (Fig. 3A). These enrichment effects are observed across the majority of PS samples, suggesting that the phenomena are common and robust (Fig. 3B). The pathways related to photosynthesis included terms such as “Photosynthetic cyanobacteria” and “Photoautotrophy” which are not surprising as there is a clear enrichment of cyanobacteria among PS-associated microbes compared to soil control samples (Fig. 2B).

**Figure 3.**
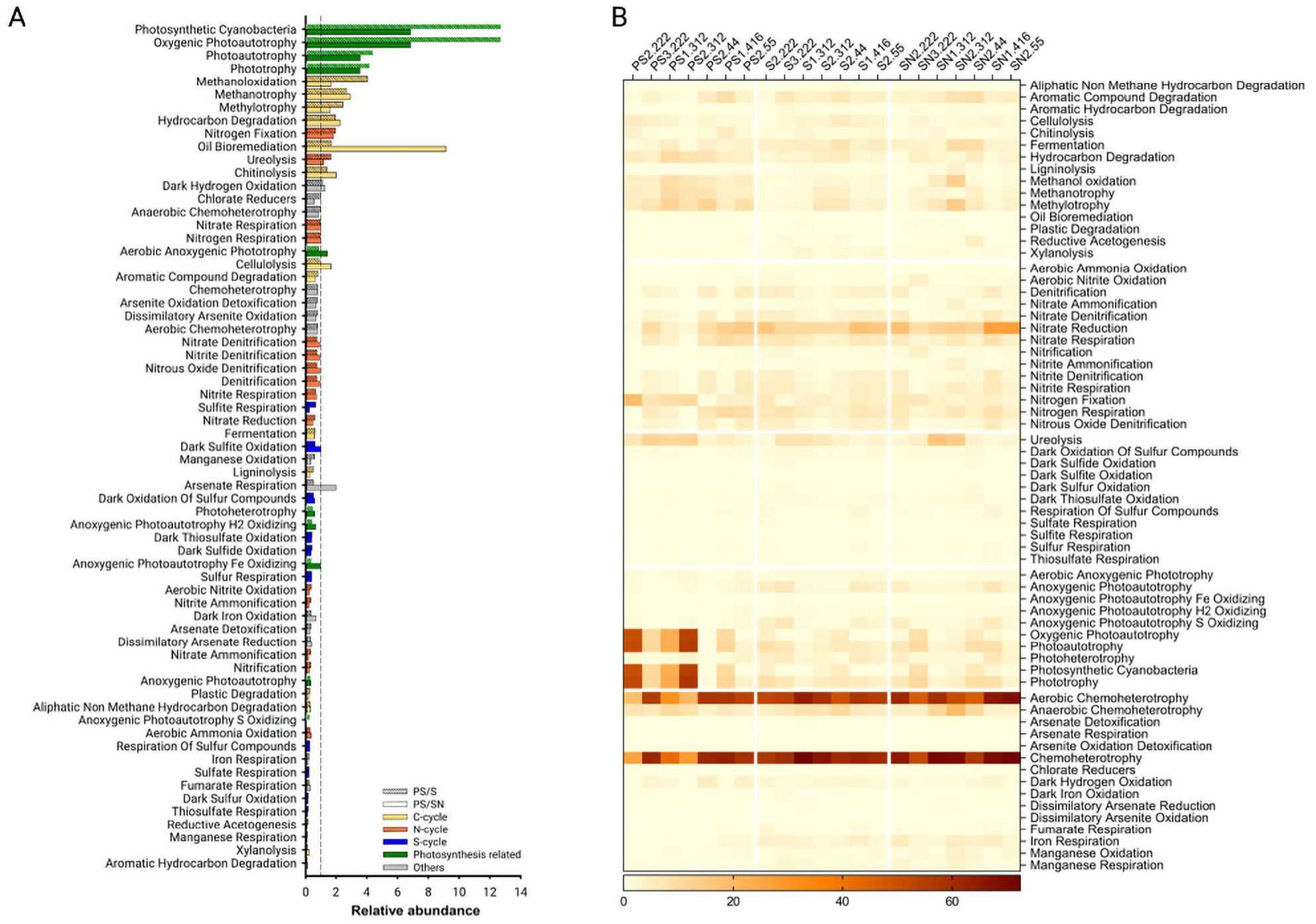
Analysis of relative abundances of metabolic pathways in PS- vs. soil-associated microbiomes. **A,** Differences in the abundances of various metabolic pathways between microbes on PS and soil samples are calculated by taking the ratios of average abundances in PS over S (solid bars) or PS over SN (hashed bars). Pathways are color-coded based on whether they are related to metabolism of carbon (yellow), nitrogen (orange), sulfur (blue), photosynthesis (green), or others (grey). **B,** Heatmap indicating the calculated abundances of each type of metabolic pathway in individual samples.

The second type of enriched pathways are those related to alkane metabolism such as “Oil bioremediation”, “Methanotrophy”, “Hydrocarbon degradation”, among others (Fig. 3A). The enrichment of these categories are not related to cyanobacteria as similar enrichment levels were observed when cyanobacteria species were removed from our sequencing results (Fig. S4B). A significant proportion of petroleum hydrocarbons are alkanes with carbon lengths of 10-100, and there is an extensive literature on bacterial strains that can degrade different chain lengths of alkanes and petroleum hydrocarbons. PS also has a carbon backbone. Thus, the enrichment of alkane metabolic pathways raises the intriguing possibility that such enzymes are involved in PS colonization or biodegradation.

### Species enriched on PS frequently encode long-chain alkane hydroxylases

The enrichment in metabolic pathways related to alkane degradation prompted us to investigate whether enzymes involved in these processes are present in species found on PS. We first sorted the species profiled in our sequencing survey based on their PS-to-soil enrichment levels (Fig. 4A, Table S3). We calculated the average enrichment across all 7 sets of samples to favor those that exhibit repeat enrichment among collection sites (Fig. 4B).

**Figure 4.**
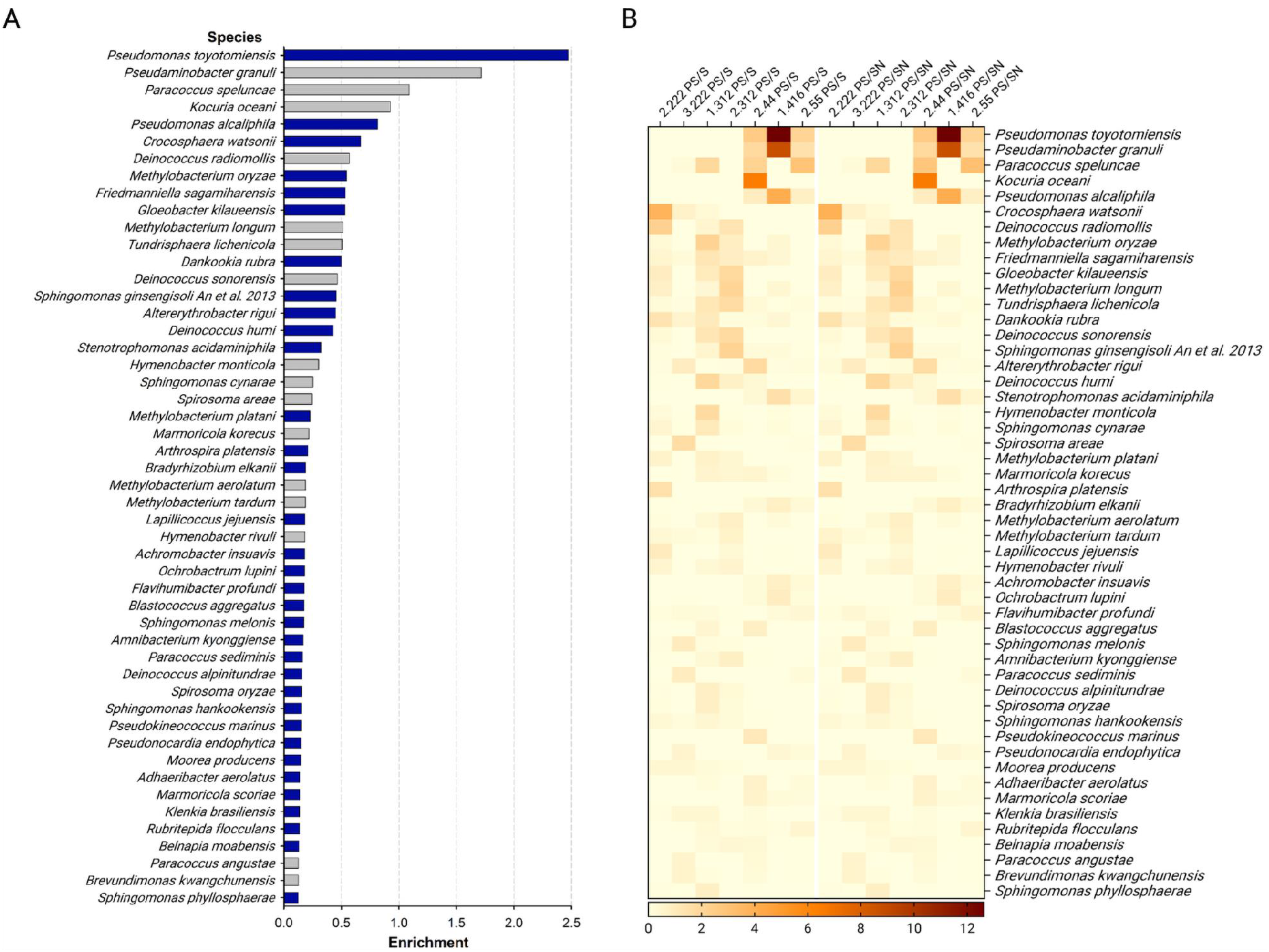
Species enrichment on PS compared to soil samples. **A,** Fold enrichment of the average presence of individual species on PS compared to the two soil samples. The bars for the species whose whole genome sequence is available are blue, those for species without are gray. **B,** Heatmap representation of individual species abundances comparing PS to S and PS to SN.

At the species level, there were not many whose fold-enrichment levels were high, but the enrichment of top candidates was clear (Fig. 4A). The species with the highest average enrichment, *Pseudomononas toyotomiensis*, is one that was first isolated from soil immersed in hydrocarbon-containing hot springs and has been reported to assimilate hydrocarbons (27). This further encouraged us to look in the genomes of the top enriched species for the presence of alkane hydroxylase homologs, and we targeted *alkB, CYP153, almA*, and *ladA* as they encode the main enzymes known to degrade hydrocarbons of various chain-lengths (18). AlkB and CYP153 typically oxidizes alkanes of less than 20 carbons whereas AlmA and LadA assimilate long-chain alkanes in the range of C_20_-C_36_ (28, 29).

For AlkB and CYP153, we found that there is modest enrichment in the top 30 most PS-enriched species compared to 30 random species profiled in our sequencing data (Fig. 5A-B, Table S4). The top and random 30 lists included only those whose whole genome sequences are available as this is a prerequisite for homolog searches. Of note, the top two species on the list, *P. toyotomiensis* and *Pseudomonas alcaliphila*, both carry *alkB* genes in their genomes (Fig. 5A).

**Figure 5.**
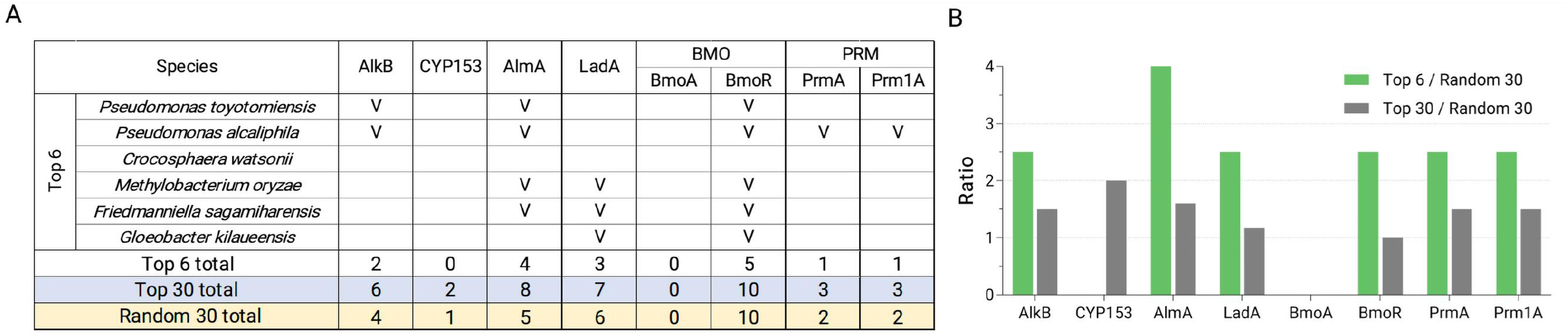
Presence of alkane hydroxylases in the genomes of top PS-enriched species and randomly chosen control species. **A,** The numbers of alkane hydroxylases found in the genomes of top PS-enriched species and random species. The numbers for the top 6 species, which are those that exhibit more than 50% PS-to-soil enrichment, are listed by individual species. **B,** Fold enrichment of the presence of alkane hydroxylases in top 6 and top 30 PS-enriched species compared to the 30 random species.

There is also mild enrichment for three of the four methane monooxygenases inspected (PrmA, Prm1A, BmoR), but the clearest enrichment observed was for long-chain alkane hydroxylases, AlmA and LadA (Fig. 5B). Of the 6 species that exhibit at least 50% enrichment on PS over soil samples, 5 encoded at least one of AlmA or LadA. The prevalence of AlmA in top 30 species compared to the random 30 list is also the highest of all alkane hydroxylases we examined (Fig. 5B). For the two most PS-enriched species, *P. toyotomiensis* and *P. alcaliphila, we* identified 3 AlmA homologs each, and for all strains of these two species whose genome sequences are available, the same 3 AlmAs are present (Fig. S5). Moreover, there is a high degree of synteny of the loci surrounding these *almA* genes; these loci are syntenic between different strains as well as between *P. toyotomiensis* and *P. alcaliphila* (Fig. S5). These results indicate that the *almA* genes are common and conserved in these two closely-related bacteria.

As in the examples of *P. toyotomiensis* and *P. alcaliphila*, many bacterial species encode more than one long-chain alkane hydroxylase in their genomes, and some carry both *almA* and *ladA* genes whereas others encode more than one AlmA or LadA proteins. Based on sequence comparisons, there appear to be multiple families of AlmA and LadA proteins (30). To investigate whether some subtypes may be more strongly associated with colonization on PS, we compared the sequences of AlmA and LadA homologs identified in the top and random species. This was done by alignment of homologs and subsequent construction of phylogenetic trees to highlight structural similarities that could delineate functional categories.

Our phylogenetic analyses revealed considerable diversity within homologs of AlmA and LadA, and there are several major subtypes for both long-chain alkane hydroxylases (Fig. 6A-B). We found that while LadA homologs from PS-enriched and random species were relatively evenly distributed across different subtypes (Fig. 6B), there are two groups of AlmA—Clans A and B—that are more highly represented in top 30 PS-enriched species with Clan B containing the densest cluster of AlmA homologs from PS-enriched species (Fig. 6A). Of the 5 top species that exhibit over 50% PS-to-soil enrichment and encode AlmA proteins, 4 had homologs in Clan B while only 1 out of 8 AlmA-encoding, nonenriched species had homologs placed in the same group. Furthermore, AlmA proteins from Clan B are also the ones with the highest similarity to the *Acinetobacter baylyi* AlmA. *Acinetobacter baylyi* is a species isolated from oil-polluted sites, and its AlmA is one of the best-characterized long-chain alkane hydroxylase (31). These again connect the biodegradation of petroleum hydrocarbons to abilities in PS colonization.

**Figure 6.**
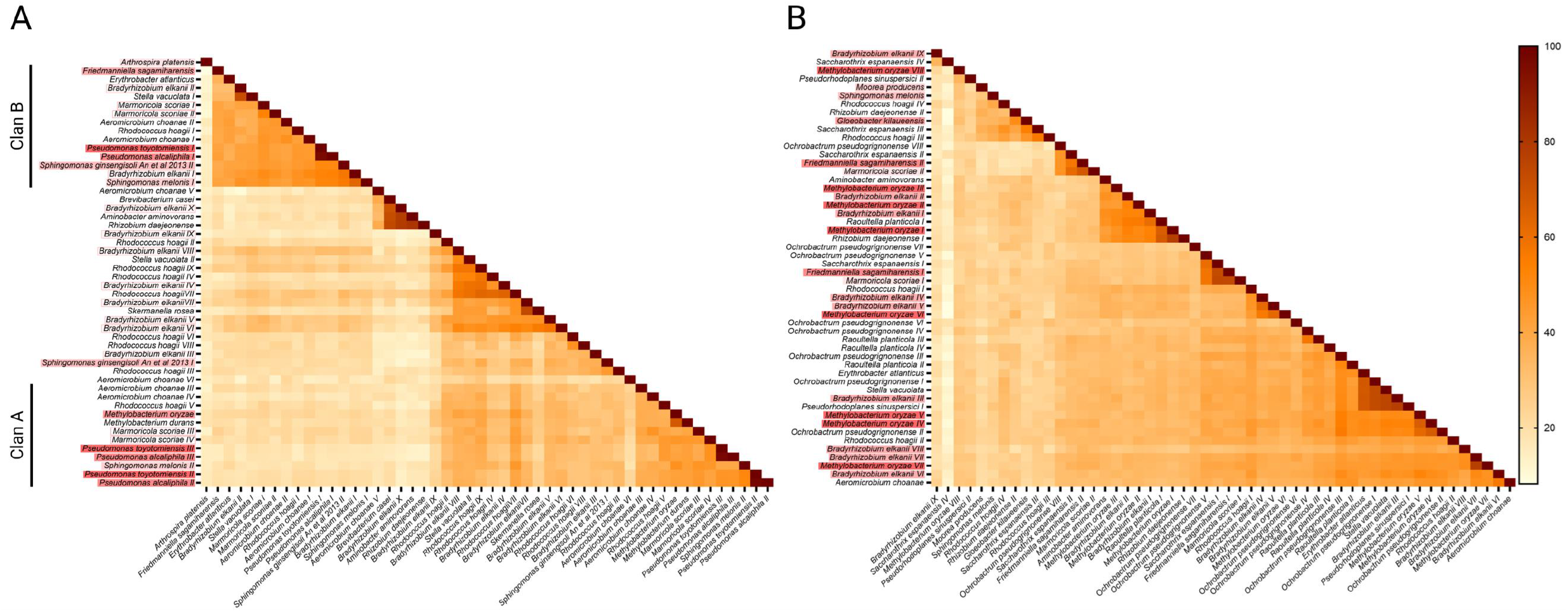
Pairwise comparisons of homologs of the two main long-chain alkane hydroxylase families, AlmA **(A)** and LadA **(B).** Percent identities between pairs are indicated as the color code on the right. Different homologs from a single species are designated with roman numerals, and the shading color of species name from dark red to none indicate strong to no enrichment of a given species on PS. In (A), two groups contain a high fraction of homologs from PS-enriched species, and they are assigned as Clans A and B.

## Discussion

Biodegradation of plastics is a great vision, but its realization still faces many significant challenges. An important starting point for the development of such technology is to identify microbial species and enzymes that have activity in plastics biodegradation so that we can start to understand and mine the potential in the microbial world for treating the plastic waste problem we face.

In this study, we profiled seven sets of PS-associated microbiome along with control soil samples near and around the site of PS collection to investigate the relationship between such plastic waste and environmental microbes. These analyses have given rise to two main insights: that select phyla are highly enriched on PS waste, and that enzymes involved in alkane hydroxylation are quite likely to contribute to bacterial colonization on PS surfaces.

The vegetation and soil pH of our collection sites were different, and these would presumably affect the microbiome in their environment, but our sequencing analysis revealed that the major microbes present in the soil of different collection sites were rather similar. In contrast, the metagenomes of the seven PS samples we obtained were much more different. The PS-enriched bacteria are not absent in the soil samples but just at much lower frequencies, suggesting that select microbes in the natural environments can gain survival advantages on the PS surface due to differences in nutrient utilization or other factors conferred by the plastic.

### Alkane degradation pathways are enriched in PS-associated microbiota

For the PS-associated microbiota, functional analysis of metabolic pathways revealed enrichment in two physiological categories: photosynthesis and alkane metabolism. The former is due to cyanobacteria being enriched on PS samples, and the latter pointed to the possibility that bacteria that encode alkane degradative enzymes are selected to grow on PS surfaces. It is very tempting to propose that certain alkane hydroxylases are reactive towards PS, thereby conferring survival advantages of microbes that secrete them on PS surfaces. However, we need to point out that the data we present here does not establish such a direct relationship.

Currently, the best-studied alkane hydroxylases are AlkB and CYP153 that process alkane with carbon chain lengths up to about 20. The presence of CYP153 in the genomes of PS-enriched species was sparse whereas AlkB is encoded by the two top species. This is a mild enrichment and could be functionally related to PS colonization especially considering that there are recent studies that reported AlkB-encoding species with the ability to biodegrade PE which also has a carbon backbone (32).

Plastics are very long polymers, thus we further examined the genomes of PS-enriched strains for the presence of long-chain alkane hydroxylases AlmA and LadA as these two are the major enzymes of this class. Quite interestingly, we found that of the species that exhibit at least 50% enrichment on PS samples, 83% encode at least one long-chain alkane hydroxylase. This is significantly higher than in the random species we screened, and it implies that long-chain alkane hydroxylases have a greater significance towards PS degradability than AlkB and CYP153.

Furthermore, the majority of top PS-enriched species contained AlmA homologs of the family that exhibits the highest similarity to the AlmA identified from the species we used as query sequence, *A. baylyi*, and this species has been shown to emulsify and assimilate petroleum hydrocarbons (33). Long-chain alkane hydroxylases are starting to be recognized as important contributors to the degradation of petroleum hydrocarbons (30, 34, 35), and there could be parallels between the species and enzymes that mediate biodegradation of petroleum and petroleum-based plastics.

### Enrichment of Cyanobacteria and Deinococcus-Thermus on PS samples

At the phylum level, the strongest enrichment we observed was for cyanobacteria. This was not due to repeat enrichment of a few cyanobacteria species in the majority of the PS samples we collected but rather the increased presence of a variety of cyanobacteria. This suggests that certain common properties of cyanobacteria grant these species growth advantages on PS. Cyanobacteria are autotrophs capable of fixing both inorganic carbon and nitrogen to organic compounds; in addition, they are known to produce a wide variety of compounds and tolerate extreme conditions. These special qualities have made cyanobacteria very valuable for agricultural and bioengineering purposes. In fact, multiple cyanobacterial strains have been reported to be able to degrade hydrocarbons (36–38), but we did not find common alkane hydroxylases (AlmA, LadA, AlkB, CYP153) to be present in the genomes of the cyanobacteria species enriched on PS.

Another interesting property of cyanobacteria is that most species produce C15-C19 alkanes for various uses related to cell membrane including maintaining cell shape and regulating cell division (39). It would not be a stretch to imagine that cyanobacteria encode mechanisms to metabolize alkanes for homeostasis purposes, but there has not been reports of such. Some studies have rather focused on non-cyanobacteria strains with hydrocarbon degradative capabilities to play the role of hydrocarbon clearance in ecosystems (40), and this could also be an explanation for why we observed enrichment of alkane hydroxylases of non-cyanobacteria species on PS surfaces.

The phylum of Deinococcus-Thermus includes many species that are extremophiles or were isolated from hazardous environments. A few of the species have been explored in various types of bioremediation processes due to their abilities to tolerate harsh or toxic environments (41–43), and our finding that this phylum is enriched on PS waste samples imply that some of their species may be particularly capable of thriving on this surface, such as being able to use PS as a source of nutrient for growth.

Bacteria in the phyla of both Cyanobacteria and Deinococcus-Thermus are ones with unique and diverse metabolism that allows them to utilize or synthesize compounds and survive in unusual environments. This is possibly why these two clades were enriched on PS samples and warrants future research to look into the metabolic properties that enable them to colonize on PS.

## Declarations

### Availability of data and material

The sequencing data in this study is deposited at the NCBI SRA under the accession number PRJNA800595.

### Author contributions

Conceptualization, S.Y.Y., S.D.F.; Data acquisition, S.W.H., Y.-H.J.; Data analysis, S.W.H., S.Y.Y.; Writing, S.Y.Y., S.W.H., S.D.F., Funding, S.Y.Y., S.D.F.

## Acknowledgements

We would like to thank the Yang lab for technical assistance. 16S rRNA gene sequencing service was provided by Biotools.

