## Supplementary Data for "Polystyrene-colonizing bacteria are enriched for long-chain alkane degradation pathways"

**Supplementary Tables**

**Table S1. Location, pH, and vegetation of collection sites.**

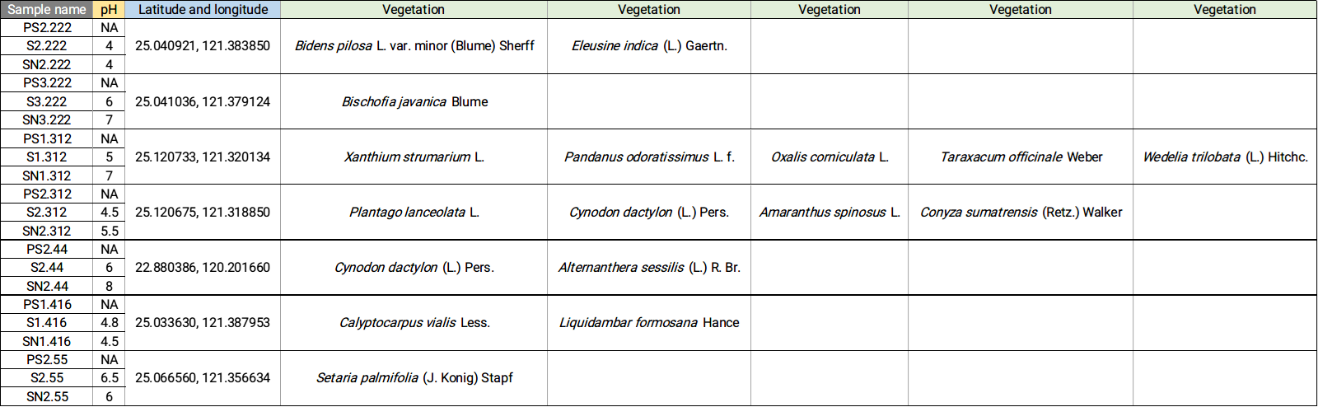

**Table S2. Relative abundances of Cyanobacteria and Deinococcus-Thermus species surveyed in the PS, S, and SN samples.**

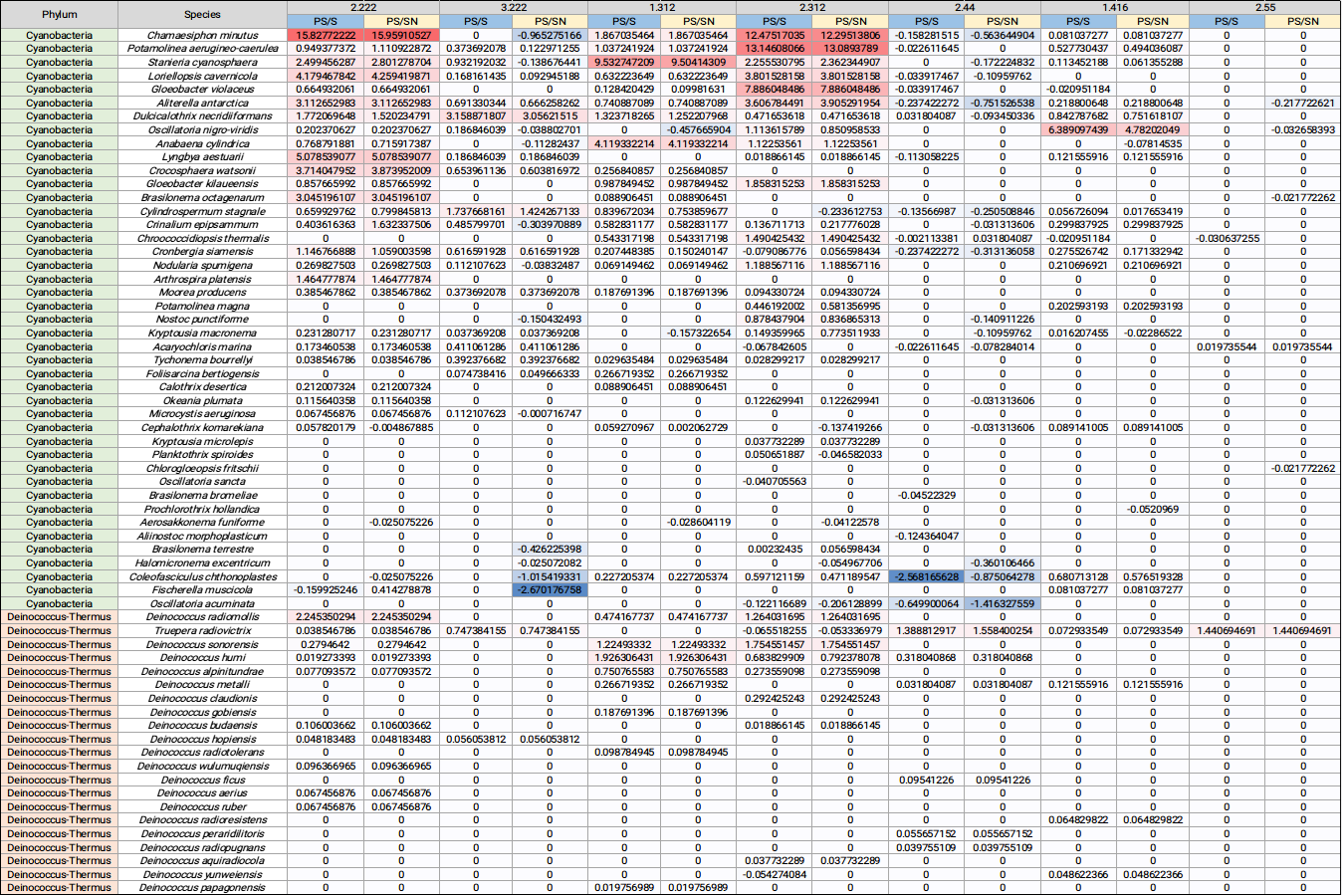

**Table S3. Top 50 PS-enriched species ranked by average levels of PS-to-soil enrichment.**

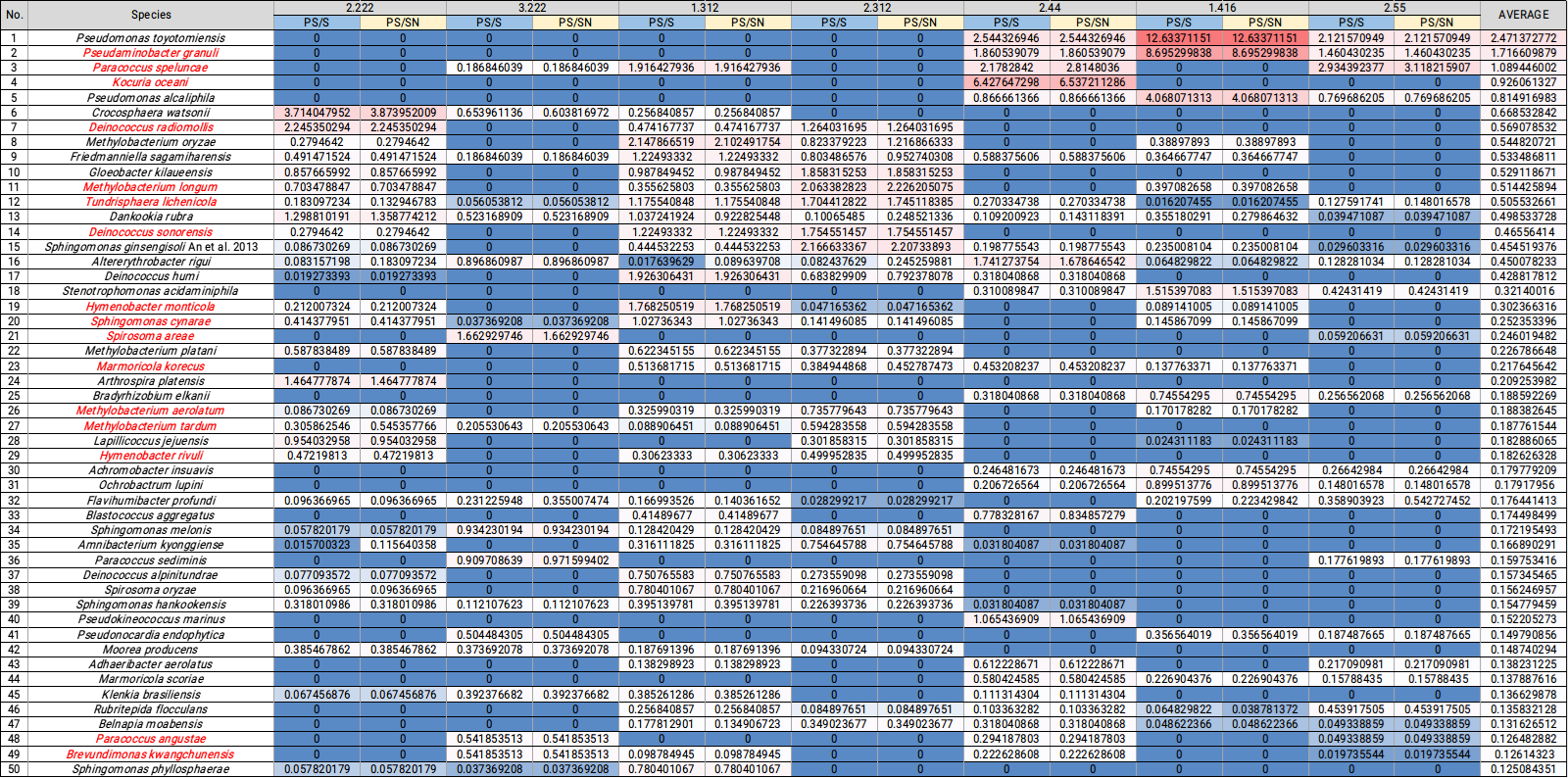

**Table S4. Lists of top 30 PS-enriched species, random 30 species, and random 60 species used in our alkane hydroxylase analysis.**

| **Top 30 PS-enriched species** | **Random 30 species** | **Additional 30 random species*** |
| --- | --- | --- |
| *Pseudomonas toyotomiensis* | *Pseudorhodoplanes sinuspersici* | *Paracraurococcus ruber* |
| *Pseudomonas alcaliphila* | *Rhizobium halotolerans* | *Oceanibaculum pacificum* |
| *Crocosphaera watsonii* | *Planctopirus limnophila* | *Paracoccus litorisediminis* |
| *Methylobacterium oryzae* | *Skermanella rosea* | *Acidothermus cellulolyticus* |
| *Friedmanniella sagamiharensis* | *Flavobacterium anhuiense* | *Fluviicola chungangensis* |
| *Gloeobacter kilaueensis* | *Rhodoligotrophos appendicifer* | *Azonexus fungiphilus* |
| *Dankookia rubra* | *Patulibacter minatonensis* | *Fonticella tunisiensis* |
| *Sphingomonas ginsengisoli An et al. 2013* | *Mycobacterium asiaticum* | *Thermogutta terrifontis* |
| *Altererythrobacter rigui* | *Desulfomonile tiedjei* | *Clostridium moniliforme* |
| *Deinococcus humi* | *Methylobacterium durans* | *Bosea robiniae* |
| *Stenotrophomonas acidaminiphila* | *Flavobacterium hercynium* | *Brevibacterium casei* |
| *Methylobacterium platani* | *Mucilaginibacter pineti* | *Acinetobacter proteolyticus* |
| *Arthrospira platensis* | *Hyphomicrobium facile* | *Legionella dresdenensis* |
| *Bradyrhizobium elkanii* | *Inquilinus limosus* | *Legionella fallonii* |
| *Lapillicoccus jejuensis* | *Metabacillus litoralis* | *Raoultella planticola* |
| *Achromobacter insuavis* | *Ochrobactrum pseudogrignonense* | *Ideonella sakaiensis* |
| *Ochrobactrum lupini* | *Duganella sacchari* | *Luteitalea pratensis* |
| *Flavihumibacter profundi* | *Erythrobacter atlanticus* | *Rhodobacter ovatus* |
| *Blastococcus aggregatus* | *Saccharothrix espanaensis* | *Edaphocola flava* |
| *Sphingomonas melonis* | *Terrabacter tumescens* | *Ornithinicoccus hortensis* |
| *Amnibacterium kyonggiense* | *Paenarthrobacter nicotinovorans* | *[Empedobacter] haloabium* |
| *Paracoccus sediminis* | *Aminobacter aminovorans* | *Hyphomicrobium hollandicum* |
| *Deinococcus alpinitundrae* | *Sphingobacterium detergens* | *Aeromicrobium choanae* |
| *Spirosoma oryzae* | *Rhizobium daejeonense* | *Stella vacuolata* |
| *Sphingomonas hankookensis* | *Edaphocola flava* | *Geobacter argillaceus* |
| *Pseudokineococcus marinus* | *Nocardioides lianchengensis* | *Rhodococcus hoagii* |
| *Pseudonocardia endophytica* | *Litoreibacter ponti* | *Streptomyces brevispora* |
| *Moorea producens* | *Labedaea rhizosphaerae* | *Micropruina glycogenica* |
| *Adhaeribacter aerolatus* | *Clostridium combesii* | *Hyalangium minutum* |
| *Marmoricola scoriae* | *Mesorhizobium tianshanense* | *Moorella thermoacetica* |

*the two random 30 lists together make up the random 60 species list

**Supplementary Figures**

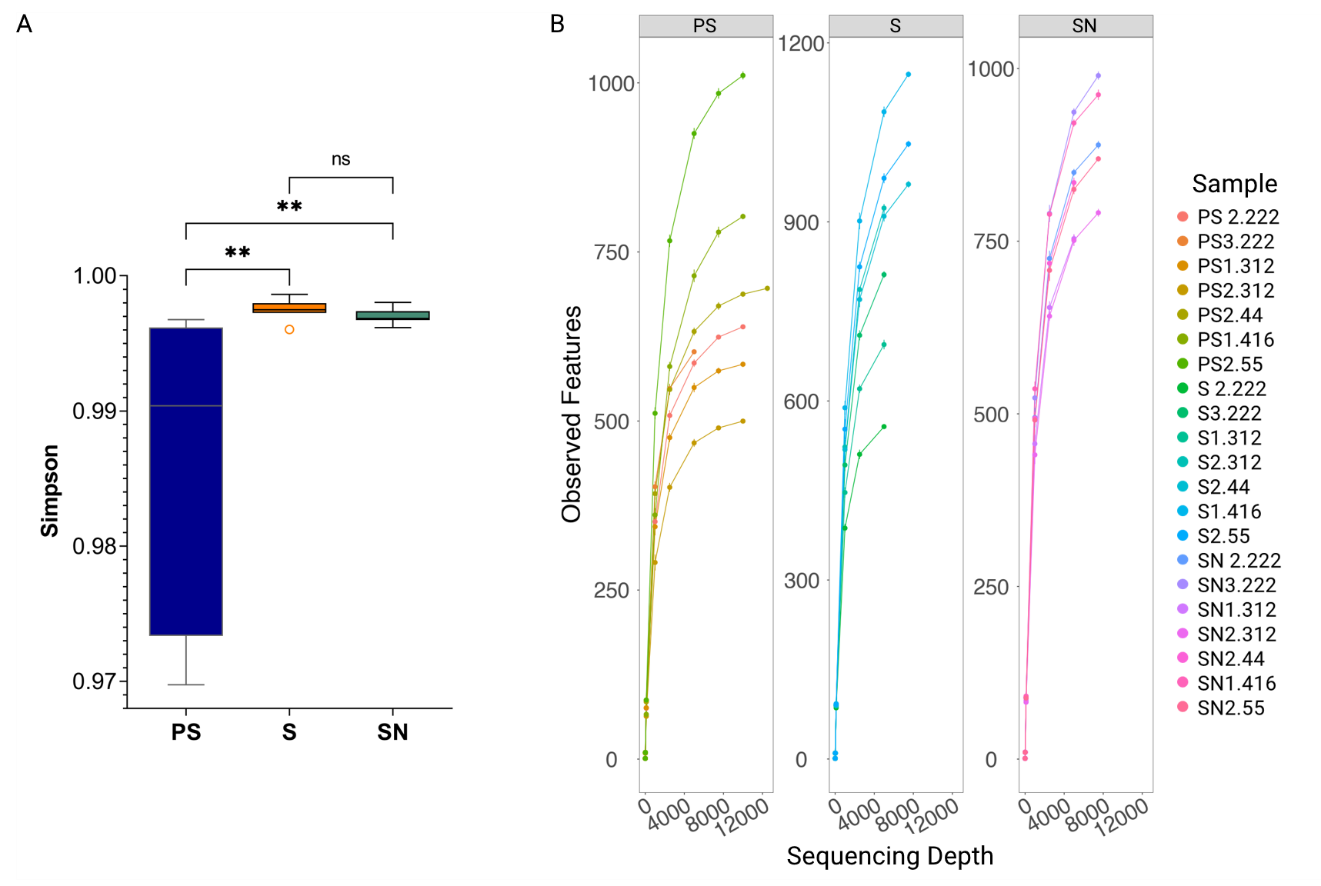
**Figure S1.** Alpha-diversity is lower in PS-associated microbiome than control soil samples. **A**, Simpson diversity indices of microbiomes associated with PS, S, and SN samples. **, P<0.05. **B**, Rarefaction curves for the PS, S, and SN samples.

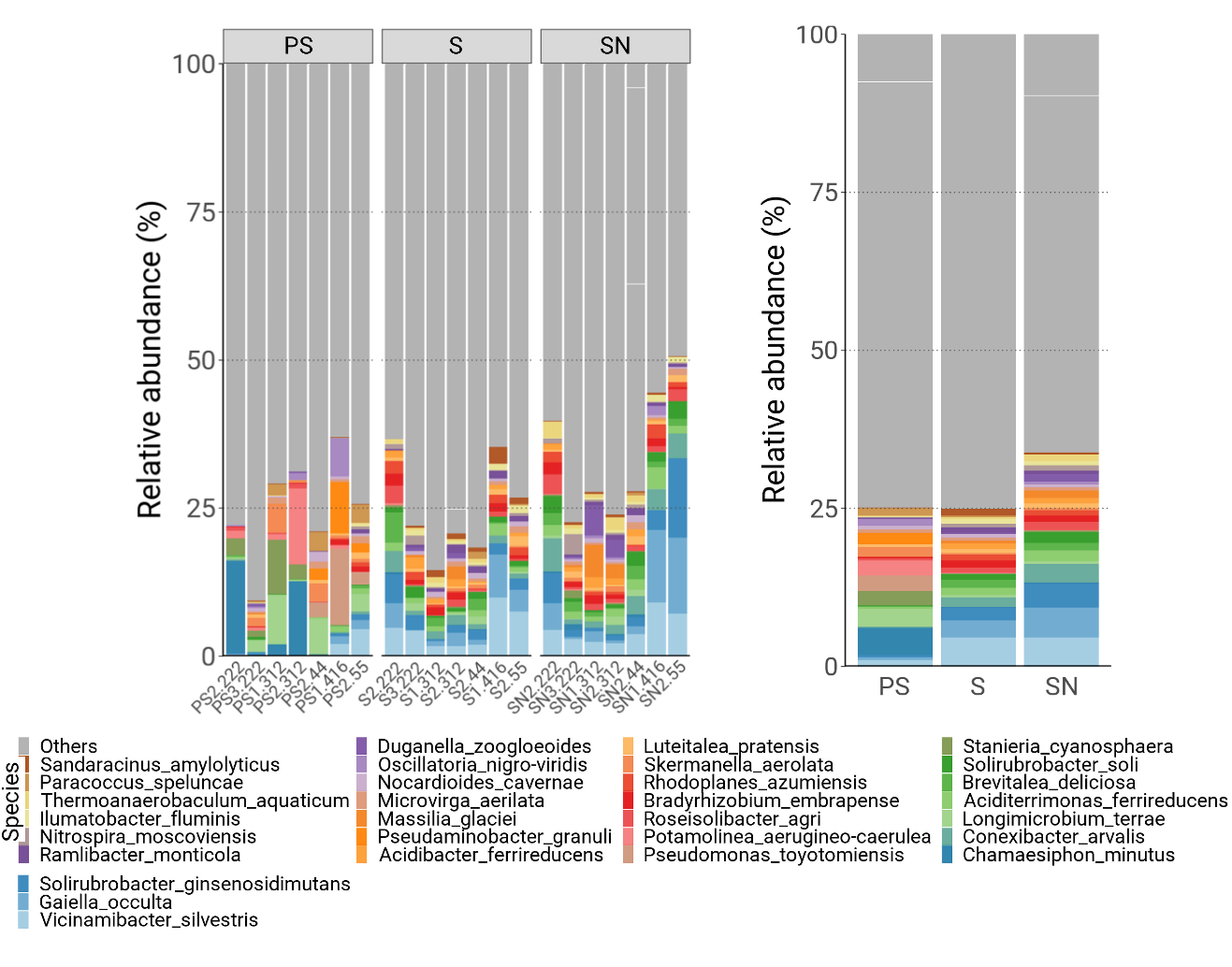

**Figure S2.** Relative abundance of the most-detected species in the PS, S, and SN samples. The bar graphs on the left depict individual plastic samples of the three categories separately whereas the graph on the right exhibits combined results of the three sample types. Color codes of the different phyla are on the bottom.

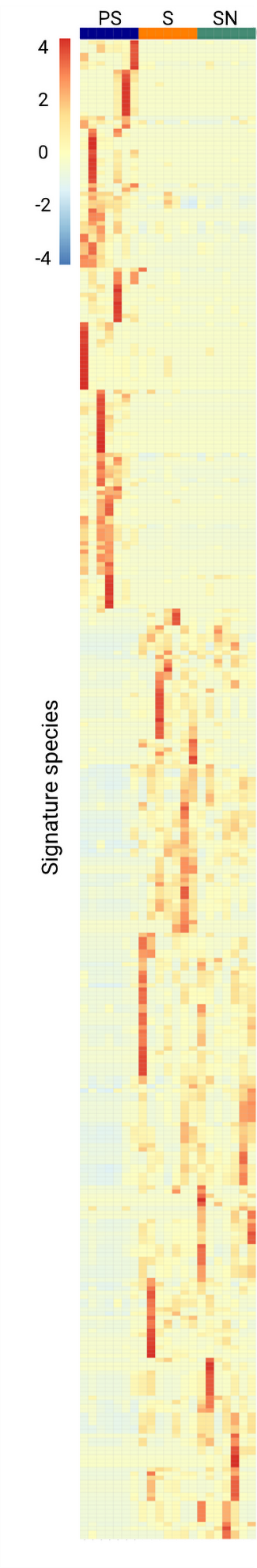
**Figure S3.** Signature species from each of the PS, S, and SN samples as calculated using the LEfSe method to demonstrate that S- and SN-associated microbes are similar to each other while dissimilar to PS-associated species.

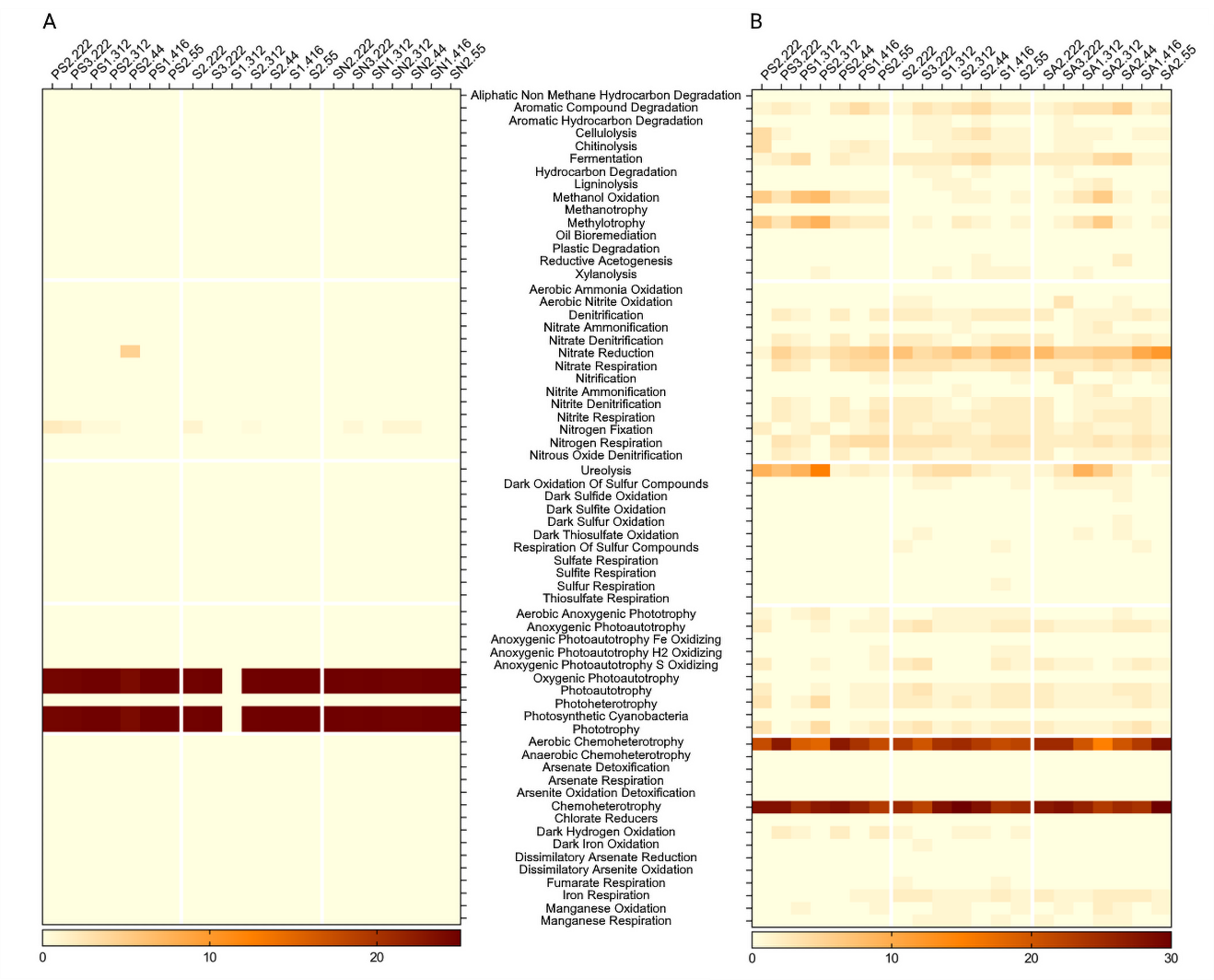

**Figure S4.** Heatmap representation of metabolic pathway enrichment analyses based on the FAPROTAX database. **A**, Analysis for the Cyanobacteria and Deinococcus-Thermus species detected in our datasets. **B**, Functional enrichment analysis of PS-associated microbes without the Cyanobacteria and Deinococcus-Thermus species detected in our datasets. The relative abundance for each pathway is indicated individually for each sample.

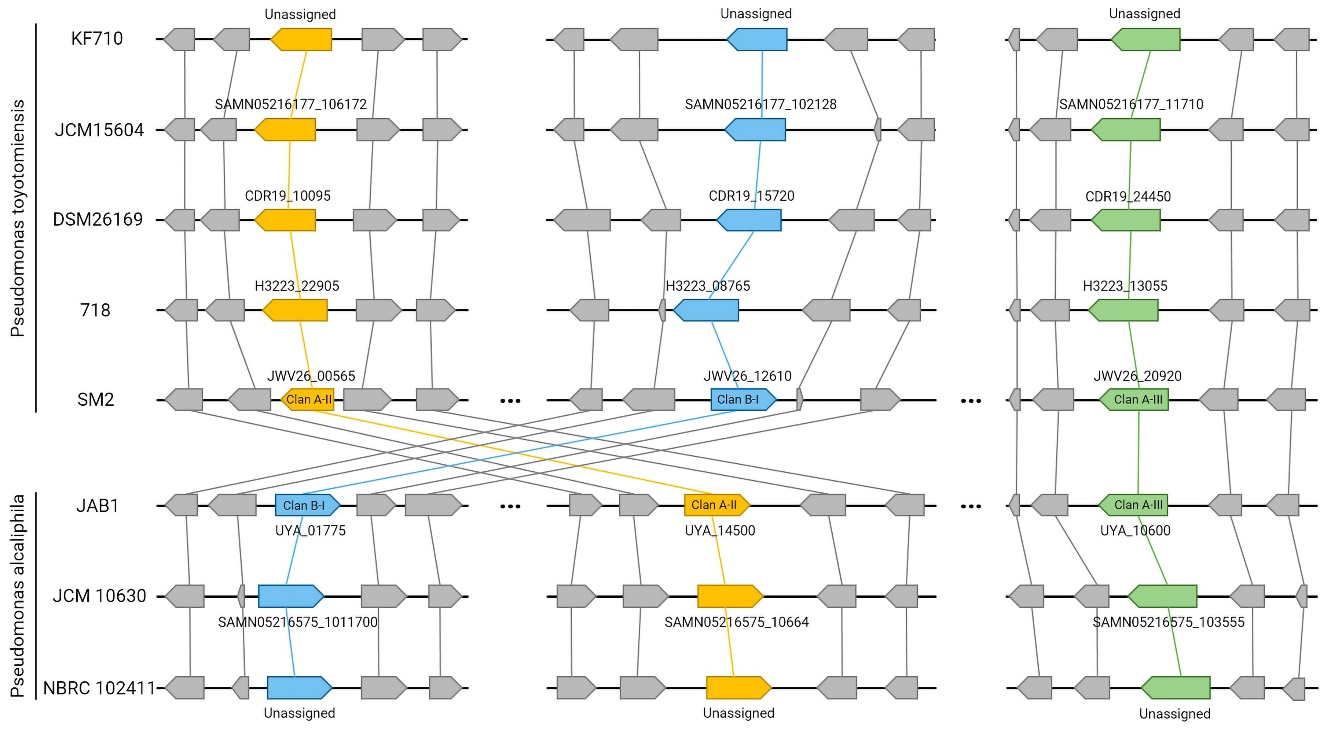

**Figure S5.** Synteny analysis of predicted genes surrounding *almA* homologs in strains of *P. toyotomiensis* and *P. alcaliphila* whose whole genome sequences are available in the NCBI database. The blue, yellow, and green boxes represent forms I, II, and III of AlmA, respectively, as categorized in Figure 6A. Strain names are indicated on the left whereas gene names for AlmA homologs are indicated near the boxes.
